# Shared population-level dynamics in monkey premotor cortex during solo action, joint action and action observation

**DOI:** 10.1101/2020.12.23.424150

**Authors:** Giovanni Pezzulo, Francesco Donnarumma, Simone Ferrari-Toniolo, Paul Cisek, Alexandra Battaglia-Mayer

## Abstract

Studies of neural population dynamics of cell activity from monkey motor areas during reaching show that it mostly represents the generation and timing of motor behavior. We compared neural dynamics in dorsal premotor cortex (PMd) during the performance of a visuomotor task executed individually or cooperatively and during an observation task. In the visuomotor conditions, monkeys applied isometric forces on a joystick to guide a visual cursor in different directions, either alone or jointly with a conspecific. In the observation condition, they observed the cursor’s motion guided by the partner. We found that in PMd neural dynamics were widely shared across action execution and observation, with cursor motion directions more accurately discriminated than task types. This suggests that PMd encodes spatial aspects irrespective of specific behavioral demands. Furthermore, our results suggest that largest components of premotor population dynamics, which have previously been suggested to reflect a transformation from planning to movement execution, may rather reflect higher cognitive-motor processes, such as the covert representation of actions and goals shared across tasks that require movement and those that do not.

**HIGHLIGHTS:**

- In PMd neural dynamics is shared across action execution and mere observation
- Task directional features are more accurately discriminated than action types
- Spatial aspects are encoded in PMd independently from specific behavioral demands
- PMd dynamics largely reflect higher cognitive-motor processes rather than strictly motor-related functions

## 1. INTRODUCTION

Classical motor-related areas are involved not just in the specification of movement kinematics and/or dynamics ^1–3^ but also in cognitive dimensions of behavioral tasks ^4–9^. Motor cortex itself is considered a fundamental node in the processing of cognitive information related to motor acts, along with other cortical and subcortical structures that contribute to motor planning and execution ^10^.

One of the most striking demostrations of the involvement of classical motor areas beyond purely motor tasks is the presence of covert representations of motor behavior without motor execution ^11^. Covert motor representations are evident in periods preceding memorized movements, when directional information is processed ^12^ and during action observation. The mirror response found originally in neurons of ventral premotor cortex in monkeys ^13–15^ expresses at single cell level the match of the neural mechanisms involved in the observation of a given action with those engaged when the observer performs the same action.

These matching operations have been shown not only during overt motor performance (e.g. when observing an agent grasping a piece of food ^14,15^) but also in more abstract contexts, involving the observation of a visual scene associated to a well-learned motor task, as when looking on a monitor a moving cursor, known to be controlled by an agent through a joystick ^16,17^. It has been proposed that, while observing sensory stimuli strongly associated to subsequent motor actions, a mental rehearsal of the motor actions occurs at the neural level, which involves operations similar to those associated to overt performance ^16,17^.

So far, the similarity of such matching operations, or the heterogeneity of context-dependent modulations of neural activity, have been mainly highlighted at a single cell level, using a traditional approach that focuses on the pattern of neural activation and its relation with movement parameters, with the exception of recent studies ^18–20^ in which the congruency of execution- and observation-related activity recorded from motor and premotor areas has been analyzed by projecting the relative neural trajectories in low-dimensional subspaces. Similarly, in this study, we adopt a dimensionality-reduction technique (Principal Component Analysis, PCA) to analyze the temporal evolution of patterns of neural activity in dorsal premotor cortex of the macaque brain, recorded during three different tasks. The first two tasks involve performing isometric actions to move a visual cursor to one of eight spatial targets arranged in a circle, in individual (SOLO) and dyadic (TOGETHER) conditions. The third task consists in the observation of the motion of a visual cursor generated by the action of another monkey (OBS-OTHER). The dimensionality-reduction approach used here permitted us to compare the temporal evolution of the system states - or trajectories in “neural space”^21–24^ - for the 3 tasks and the 8 different spatial targets. Notably, while other studies measured neural population dynamics during one’s own action planning and execution ^25^, our experimental paradigm allowed comparing them with population dynamics during an action observation task that does not require action generation.

We addressed two main open questions. First, we asked how task-related variables (e.g., movement initiation, direction of cursor’s motion, task identity) are coded at the population level in monkey premotor cortex. Second, we asked whether the same population-level coding is shared across tasks having different behavioral demands - and especially across tasks that require (SOLO and TOGETHER) or do not require (OBS-OTHER) overt action generation. Henceforth, we will refer to SOLO and TOGETHER tasks as “motor tasks” and OBS-OTHER as a “non-motor” task, for brevity.

## 2. MATERIALS AND METHODS

### 2.1 Animals and tasks

Two adult male rhesus monkeys (*Macaca mulatta; monkey S* and *monkey K*; body weight 7.5 and 8.5 Kg, respectively) were trained as a pair to perform an isometric hand force center-out task, in two different conditions: 1) individually (SOLO) or ii) in a joint-action context, where each animal had to coordinate its force with its companion to achieve a common goal (TOGETHER). During the recording sessions, the two animals sat next to each other (Fig. 1B), in front of a 40-inch monitor at a distance of 150 cm from the eyes. Experimental and surgical procedures were performed in conformity with European Directive (63-2010 EU) and Italian (D.L. 26-2014) laws on the use of nonhuman primates in scientific research, and under authorization of the Ministry of Health of Italy to Alexandra Battaglia-Mayer. During the experimental procedures, all efforts were made to minimize animal suffering. Details about surgical procedures have been reported previously ^26^.

**Figure 1.**
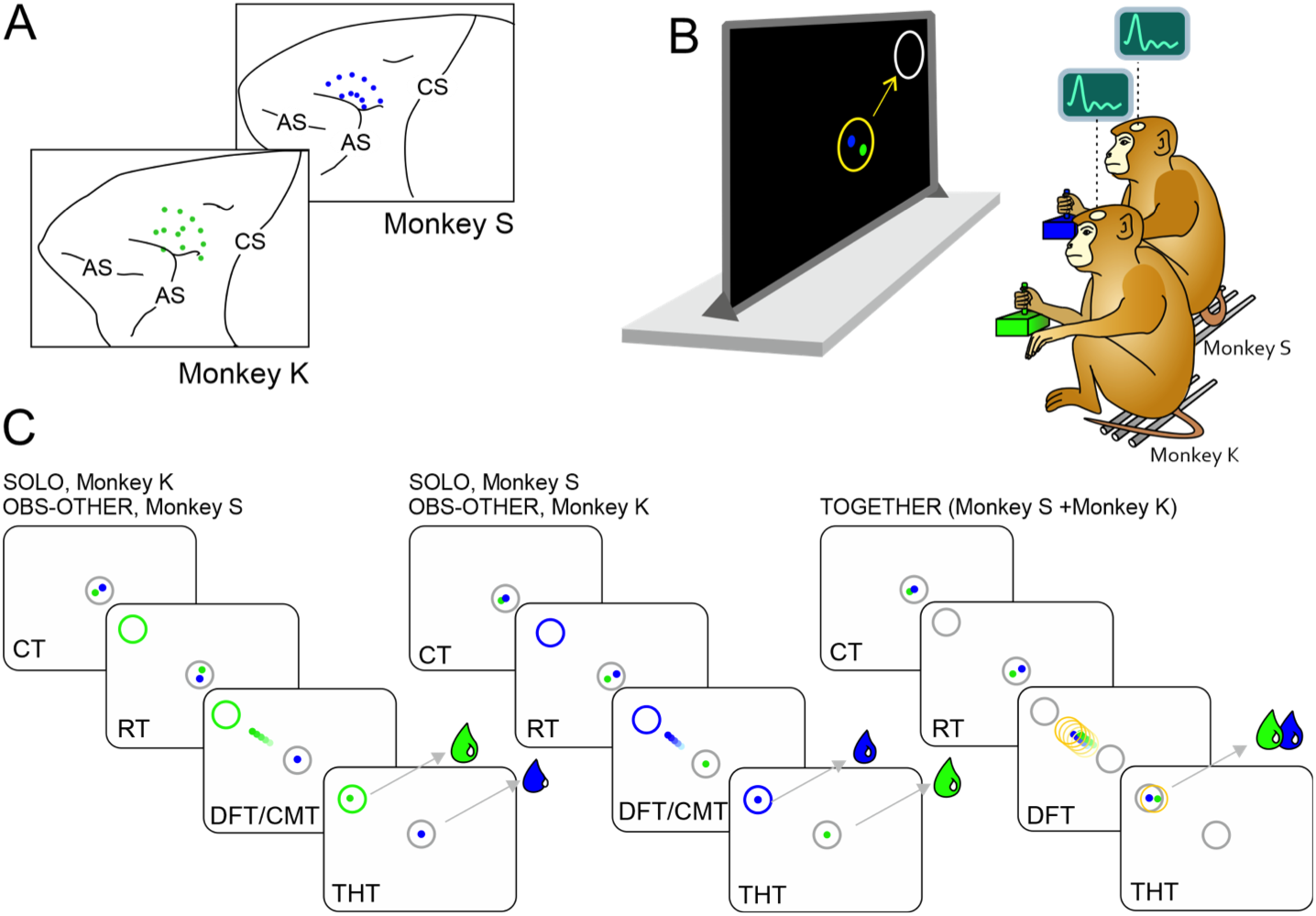
Recording sites, experimental apparatus and tasks. ***A.*** Recording sites in areas F7/F2 of Monkeys S and K. ***B.*** Monkeys S and K sat next to each other and controlled through an isometric joystick their own cursors (S, blue dot; K, green dot) displayed on a screen. ***C.*** At the beginning of each trial, the monkeys were required to bring their cursors from an offset position (not shown) to the central white circle (grey in C) for a variable Control Time (CT). In the SOLO task, each monkey, instructed by the color of one peripheral target (S, blue; K, green) presented in one of 8 potential locations, had to bring its cursor on this target, in a subjective reaction time (RT), by applying individually a hand force pulse on the joystick (Dynamic Force Time, DFT) to get a liquid reward. During the SOLO trial of one monkey, the other animal observed the moving cursor controlled by its mate (OBS-OTHER task) and was only required to actively maintain its cursor within the central circle until the end of the trial, to obtain its reward dose. In the SOLO and OBS-OTHER trials, the reward was delivered to each animal based on its individual (successful) performance, independently of the partner’ behavior. In the TOGETHER trial, instructed by the white color of the peripheral target (***B***, grey in ***C***), both monkeys had to act jointly, by coordinating their force output in direction and timing, to guide together a common yellow circle from the center to the peripheral target to get both their reward doses. For the successful performance of the task, the animals had to keep their own cursors within a maximal inter-cursor distance for the entire DFT)application and until the end of the Target Holding Time (THT). Lack of inter-subject coordination resulted in unsuccessful trials and neither animal was rewarded. Notice that the DFT of the SOLO trials of one monkey corresponds to a cursor’s motion time (CMT) interval of the OBS-OTHER task of the other animal.

Details about the behavioral tasks have been already reported in ^26,27^. Briefly, each monkey applied a hand force on an isometric joystick (ATI Industrial Automation, Apex NC), to control and move a visual cursor in a center-out task, from a central position toward on one of the 8 peripheral targets, placed at an eccentricity of 8 degrees if visual angle (DVA). The joystick consisted of a force-transducer which measured the forces in two dimensions (Fx, Fy), in absence of any hand/arm displacement. Animals had to move the cursor from the central to the peripheral target by applying a monotonically increasing isometric force ramp to the joystick in the appropriate direction, with the (x,y) coordinates of the cursor linearly related to *F_x_* and *F_y_* components of the applied force,i. e. 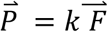 where 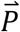 is the position of the cursor with respect to the central position and 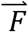 the force applied by the animal on the joystick.

To bring the cursor from the center to the peripheral targets a force pulse of about 3.2 N was required. Each monkey was trained to control its own colored cursor (blue for *Monkey S* and green for *Monkey K*; diameter 0.6 DVA. The neural activity was recorded when the animals performed the isometric task under two different conditions (SOLO and TOGETHER trials), as well as when they observed (OBS-OTHER trials) the motion of the visual cursor controlled by the other animal (namely, observing the SOLO trials of the other monkey), without any dynamic force application. During the experiment, the animals used the right arm, contralateral to the recording chamber, while their left arm was gently restrained.

In the isometric SOLO and TOGETHER tasks (Fig. 1C), trials started with the appearance of a central white circle. The animals had to bring their own cursors within it from an offset position, keeping them there for a variable control time (CT, 500-600 ms), by exerting a small static hand force. Then, a peripheral target (outlined circle, 2 DVA in diameter) appeared in one of eight possible locations. The color of the target circle indicated which of the two animal had to move the cursor (blue: SOLO for monkey S; green: SOLO for monkey K, white: TOGETHER, both monkeys), hence the type of task (SOLO, TOGETHER and OBS-OTHER) to be performed (Fig. 1B). In the SOLO trials, one monkey performed the task individually, by applying a force pulse, after a subjective reaction-time (RT), to bring its own cursor from the center toward the peripheral target (Dynamic Force Time, DFT; Fig. 1C), to obtain a liquid reward. When one animal moved its cursor in a SOLO condition, its companion had to keep its own cursor inside the central target, in order to get a liquid reward, until the end of the SOLO trial of the other monkey. During this DFT time, the companion animal observed the result of its partner’s action consisting in the motion of a cursor on the screen (cursor’s motion time, CMT). Therefore, from the perspective of this animal, these trials were those defining the ‘OBS-OTHER’ condition (Fig. 1C). Hence, the DFT of the SOLO trials of one monkey coincides with a cursor’s motion time (CMT) interval of the OBS-OTHER task of the other animal. Note that this setup is different from other action observation studies. In fact, the isometric nature of the task did not provide the acting monkey and neither its observing partner any visual cue about the performed action, other than the resulting motion of a visual cursor on the screen. In both SOLO and OBS-OTHER trials, the reward delivery to each animal depended only on the success of its own performance, which was independent from the partner’s behavior. Although controlled (see below), no constraints were imposed on eye movements, since under this condition we were interested in studying the natural oculomotor behavior of each animal.

Besides the SOLO condition, the isometric force application was tested also in the interactive context of the TOGETHER task. When the peripheral target was white in color, both monkeys were required to coordinate their force output in space and time to bring together a common visual object (yellow circle) (Fig. 1C), toward it. The yellow circle appeared simultaneously to the peripheral target in the center of the workspace. To succeed in their common goal and to prevent aborting the trial, the monkeys had to maintain a maximum inter-cursor distance limit, coinciding with the diameter of the yellow circle (IDmax, 5 degrees VA). The moving yellow circle controlled by the two animals was centered at any instant at the midpoint of the two cursors, and once it reached successfully its final location, both monkeys received simultaneously an equal amount of liquid reward. The amount of reward dispensed was identical across task conditions.

Twenty-four trial types were collected, consisting of two SOLO conditions (one for each animal, which corresponded to the OBS-OTHER condition for the non-acting monkey) and one TOGETHER condition, all executed for eight different peripheral target locations (3 conditions x 8 directions). These 24 trials were presented in an intermingled fashion design and pseudo-randomized within each replication, until each of them was performed succesfully. Once a replication of 24 trials was completed, trials belonging to the successive replication were presented in a new pseudo-random order. A minimum of 8 replications were performed by the animals, leading to a minimum set of 192 trials (3 conditions x 8 directions x 8 replications), defining one block.

Finally, a saccadic eye movement task (EYE) was used as a control condition to evaluate the influence of eye-related signals on the neural activity recorded during the other task types. In separate blocks of trials, each animal was required to fixate initially a central white square for a variable CT (700-1000 ms), and then to make a saccade toward one of 8 peripheral targets (8 DVA eccentrity), which appeared at the completion of CT. Eye position and movement were recorded through a non invasive infrared oculometer (Arrington Research).

### 2.2 Electrophysiological recordings

Single-unit activity was recorded from dorsal premotor cortex (PMd; area F7/F2; area 6; Fig. 1A) of the two co-acting monkeys. Neurophysiological recording performed during the above-mentioned tasks resulted for each monkey in three sets of data, relative to the three tasks in which the animal behavior was tested (SOLO, TOGETHER, OBS-OTHER). The neural activity was recorded extracellularly using two separate 5-channel multiple-electrode arrays (Thomas Recording, Giessen, Germany) from the two brains. The activity of each unit was collected in blocks of 192 trials (corresponding to 8 directions x 3 task types x 8 replications). In each session (1-3 session/day), constituted by the block of 192 trials, we recorded on average 6.8±3.7 units/session in monkey S, and 8.6±4.3 units/session in monkey K, for a total of 37 experimental sessions.

### 2.3 Data analysis

#### Cell modulation and directional tuning

For each neuron, the modulation of its activity was evaluated in two separate behavioral epochs of interest, i.e. RT and DFT, through a one way ANOVA (factor, ‘epoch’, p<0.05). A cell was defined as being modulated in a given epoch if the firing rate measured during the RT (or DFT/CMT) was significantly different from that of the control time (CT). Another one-way ANOVA was used to analyse significant differences of the firing frequencies, in a given epoch across the three task conditions (SOLO, TOGETHER, OBS-OTHER), followed by Bonferroni test for post hoc comparison).

To assess that a cell was “directionally-tuned” we used the same method adopted in ^26^, consisting of a bootstrap procedure ^28^ aimed at evaluating the statistical significance of its directional tuning. The bootstrap was applied to the tuning strength (*TS*), defined as the amplitude of the mean vector expressing the firing rate in polar coordinates ^29^. A shuffling procedure randomly reassigned single-trial data to different target directions for 1000 times and the *TS* from the shuffled data was determined. A cell was labeled as “directionally-tuned” in a specific epoch of one task, if the *TS* value calculated from the original unshuffled data was higher than the computed confidence limit (p<0.05).

#### Neural space analysis

We conducted a principal component analysis (PCA) – also called “neural space” analysis ^30–32^ – on the the neural activity of a population of 384 premotor cells (200 neurons recorded in monkey K, 184 neurons in monkey S), obtained from a larger dataset of 471 neurons, whose activity have been analysed in ^26^. The cells selected for the present study were those for which the activity was collected in at least 8 replications in all the 3 task types, in all the 8 cursor’s motion directions. For example data from single monkeys, see Fig. S6.

Our analysis considered 192 trials: 3 task types by 8 directions by 8 repetitions. We first averaged the firing rate for each neuron across the 8 repetitions. Then, we submitted the resulting 24 averaged firing rates from each neuron to 2 separate PCAs, one aligned to target onset and one aligned to cursor’s motion onset. Finally, we projected the 192 trials (aggregated by task types or directions, see below) onto the first 4 PCs of each PCA.

The analysis was performed in two different time intervals, corresponding to different activity alignments: i) the interval spanning from 200 ms before (TgOn; 0 ms) to 400 ms after target onset; ii) the interval from 500 ms before (CMon) to 200 ms after cursor’s motion onset. In both instances we adopted a 20 ms time binning. It is worth noticing that in our task the peripheral target onset was informative both about the direction of the cursor’s motion and the type of action to be performed (SOLO, TOGETHER, OBS-OTHER).

To assess the robustness of the population activity during different tasks and cursor’s motion directions, we performed a bootstrap test for each neural space trajectory ^33^. The bootstrap procedure consisted of a random resampling with replacement of the firing rate data of each cell 10,000 times in each (20ms) time bin and for each of 24 (3 task types x 8 directions) block conditions. Then, we projected these data on the first 4 PCs (X1-X4) and grouped them by tasks and directions, in order to produce a distribution of the difference in the mean trajectory at each data point. The resulting confidence intervals showed if and when in time the compared distributions were significantly different. Therefore, all the results shown in Figs. 3–6 obtained with a bootstrap procedure are such that the non-overlapping portions of neural space trajectories lies in the 5%–95% percentiles of the distribution of resampled differences. Consequently, they can be considered significantly distinct and discriminable at p < 0.05.

To compare quantitatively the differences between the encoded variables (such as direction and task) we considered a 4D neural space, formed by the first 4 components X1-X4 emerging from the PCA (or 12D neural space in the Supplementary materials). We adopted a distance metric between neural trajectories in this 4D space, using the Root Mean Square (RMS) of the Euclidean distance computed at each 20 ms time bin. In particular, given two neural trajectories **p**(*t*) and **q**(*t*), at each 20 ms time bin *t*, we computed the K=4 coefficients of each trajectory projected into the 4D space, i.e., c^**p**^(t) = cp_1_(t), … cp_k_(t) and c^**q**^(t) = cq_1_(t), … cq_k_(t), respectively. Thus, at each time step, we did define the quantity:

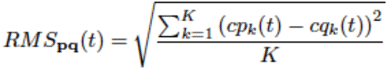

Note that the RMS distance allows comparing different projections on different space dimensions, as it is normalized for the number of components. When we consider a number N (in our case N=10,000) of bootstrap samples, we can straightforwardly compute for each time step a distance between sets of samples, **p** ∈ P, **q** ∈ Q (e.g., samples belonging to different tasks and directions), by means of the following equation:

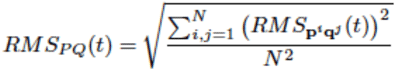

Our analysis using RMS distance was based on 24 trajectories (3 tasks x 8 directions) in 4D neural space, with alignment on target onset (Fig. 5).

## 3. RESULTS

A principal component analyses (PCA) – or “neural space” analysis ^30–32^ – was conducted on the neural activity of a population of 384 premotor cells, obtained from a larger dataset of 471 neurons from dorsal premotor cortex (PMd; areas F7/F2; part of area 6; Fig. 1A), recorded from two monkeys (204 neurons in monkey S; 267 neurons in monkey K), during the execution of different visuomotor tasks. Within the same sessions, the monkeys performed two isometric hand force tasks in different contexts (SOLO and TOGETHER) and an observation task in absence of dynamic force production (see Fig. 1 and Methods).

The single-unit activity of the 471 cells recorded under the same task conditions was analysed in a previous study ^26^, where a detailed description of the behavioral tasks is reported. In brief, two monkeys were sitting in front of a large monitor (Fig. 1B), and each held in its hand an isometric joystick to control its own visual cursor (blue for monkey S; green for monkey K). In the isometric tasks (Fig. 1C), the animals were instructed, by the color of a peripheral target, whether to bring their cursors from the central position to that target, individually (SOLO, blue or green target; Fig. 1C), or acting jointly with their companion (TOGETHER, white target; Fig. 1C). The latter condition required coordinating their forces to guide a common object (yellow circle) from the center to its final target location. During the TOGETHER condition, the instantaneous position of the moving yellow circle coincided with the midpoint of the coordinates of the two cursors, controlled simultaneously by the two animals. During the execution of the SOLO trials of one monkey, the partner was required to statically maintain its own cursor in the central position, without any dynamic hand force application, while observing on the screen the cursor’s motion controlled by its partner (OBS-OTHER trials), (Fig. 1C). Therefore, the neural activity recorded from each animal was studied during the performance of three different tasks, SOLO, TOGETHER and OBS-OTHER.

### Single cell analysis

The animal behavior in the three different tasks is described in detail in previous studies ^26,27^. As already reported ^26^, 66% (311/471) of our original dataset of cells were modulated at least in one of the three tasks, and 39% (186/471) in more than one task.

As expected, the spatial structure of the task highlighted the directional nature of the neural activity in all task types (Fig. 2A-B). Please note that another study using the same setting shows that this directionality is not influenced by the fact that the two monkeys were side-by-side (e.g., no left-right directional bias was found) ^34^. In Fig. 2A we show an example of neural activity of a single cell which was significantly modulated during the DFT in both SOLO (one-way ANOVA, F(1,127)=[20.5], p=1.36e-05) and TOGETHER (one-way ANOVA, F(1,127)=[36.7], p=1.48e-08) tasks, even though differently (one-way ANOVA, F(2,191)=[47.69], p= 1.70-17, followed by the Bonferroni post hoc test). In Fig. 2B we show another neuron active during the OBS-OTHER task (CMT; one-way ANOVA, F(1,159)=[130.7], p=1.93e-22), and not during the DFT of the two isometric tasks. For each cell, we report in the form of rasters and spike density functions the activity recorded in the three task types (SOLO, green; OBS-OTHER, blue; TOGETHER, red), in the 8 directions of cursor’s motion.

**Figure 2.**
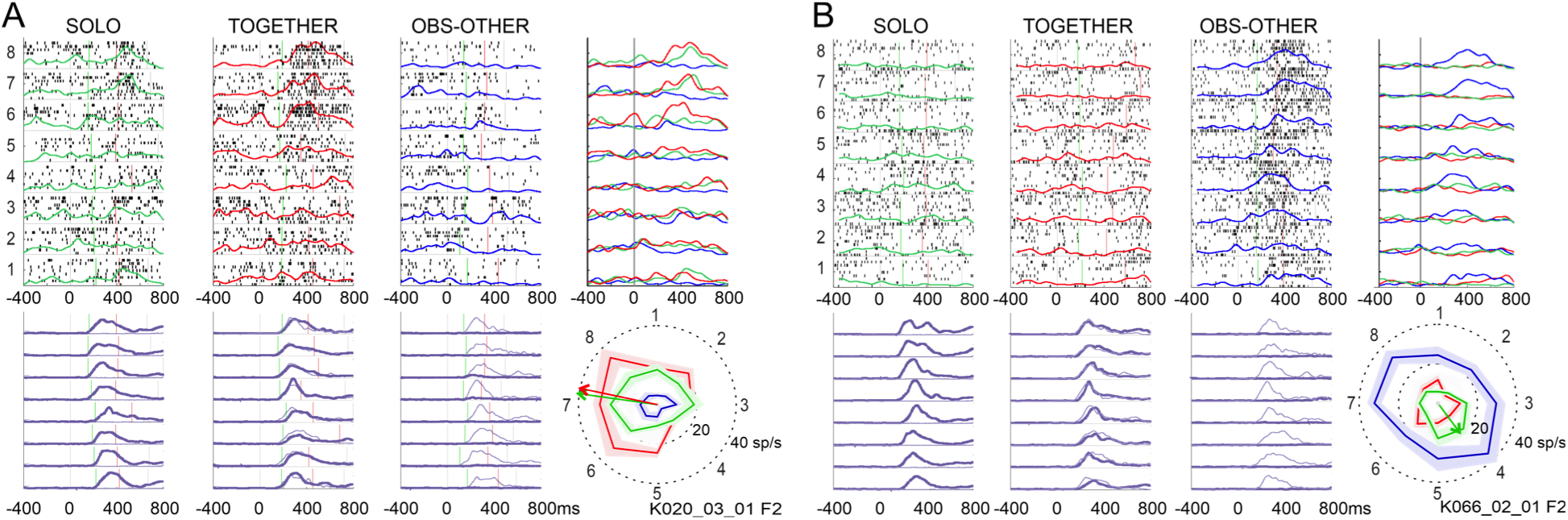
Examples of cell activity recorded in premotor cortex. ***A.*** Neural activity of a cell modulated during the isometric force application. The cell is active, during the DFT in both SOLO (one-way ANOVA, F(1,127)=[20.5], p=1.36e-05) and TOGETHER (one-way ANOVA, F(1,127) = [36.7], p=1.48e-08) tasks, even though differently (one-way ANOVA, F(2,191)=[47.69], = 1.70-17, followed by the Bonferroni post hoc test), during both the SOLO and TOGETHER tasks, and not during the OBS-OTHER trials. (***B***) Activity of another cell modulated during the OBS-OTHER task (CMT; one-way ANOVA, F(1,159) = [130.7], p=1.93e-22), and not during the DFT of the two isometric tasks. In ***A,B*** the neural activity is plotted for directions 1 to 8 in the form of raster plots, where each dot represents an action potential, as well as spike desity functions for each direction. The spike density functions calculated for the three tasks are overlapped (SOLO, green; TOGETHER, red; OBS-OTHER, blue) for direct comparison. The activity is also reported in the form of polar plots, where the mean firing rate computed during the DFT (for SOLO and TOGETHER) or the corresponding epoch (CMT) for the OBS-OTHER trials is plotted in the 8 directions, to highlight the directional nature of neural activity. Please note that for better readability, the data points in the polar plots are interpolated using arbitrary (here, straight) lines. For each cell, the animal’s behavior is reported by showing, below the raster-plots, the corresponding speed profiles (purple curves) of the cursor guided by the monkey from which the spiking activity was recorded (thick curve) or by its partner (thin curve). In all panels, 0 ms corresponds to target onset, while green and vertical bars indicate cursor’s motion onset and the time of its arrival on the final location, respectively.

### 3.2 Neural space analysis: first four principal components

We conducted two PCA analyses, on neural data aligned to target onset and to cursor’s motion onset, respectively. For each analysis, we identified the four highest-ranked principal components X1-X4 (i.e., those explaining more variance), which captured 26.1% of the variance when aligned on target onset, and 34% when aligned on cursor motion onset. We aggregated their activity according to the three tasks (i.e., SOLO, OBS-OTHER and TOGETHER; Fig. 3) or to the eight directions of cursor’s motion (Dir 1-8; Fig. 4). Note that in the two motor tasks (SOLO and TOGETHER) “direction” may refer to both the direction of isometric force output and the direction of cursor’s motion, but in the non-motor task (OBS-OTHER) only the latter applies. Here, we use “direction” to refer to the direction of cursor’s motion, which applies to all the tasks. In both Figs. 3 and 4, panels A and B show the alignment to target onset and cursor’s motion onset, respectively.

**Figure 3.**
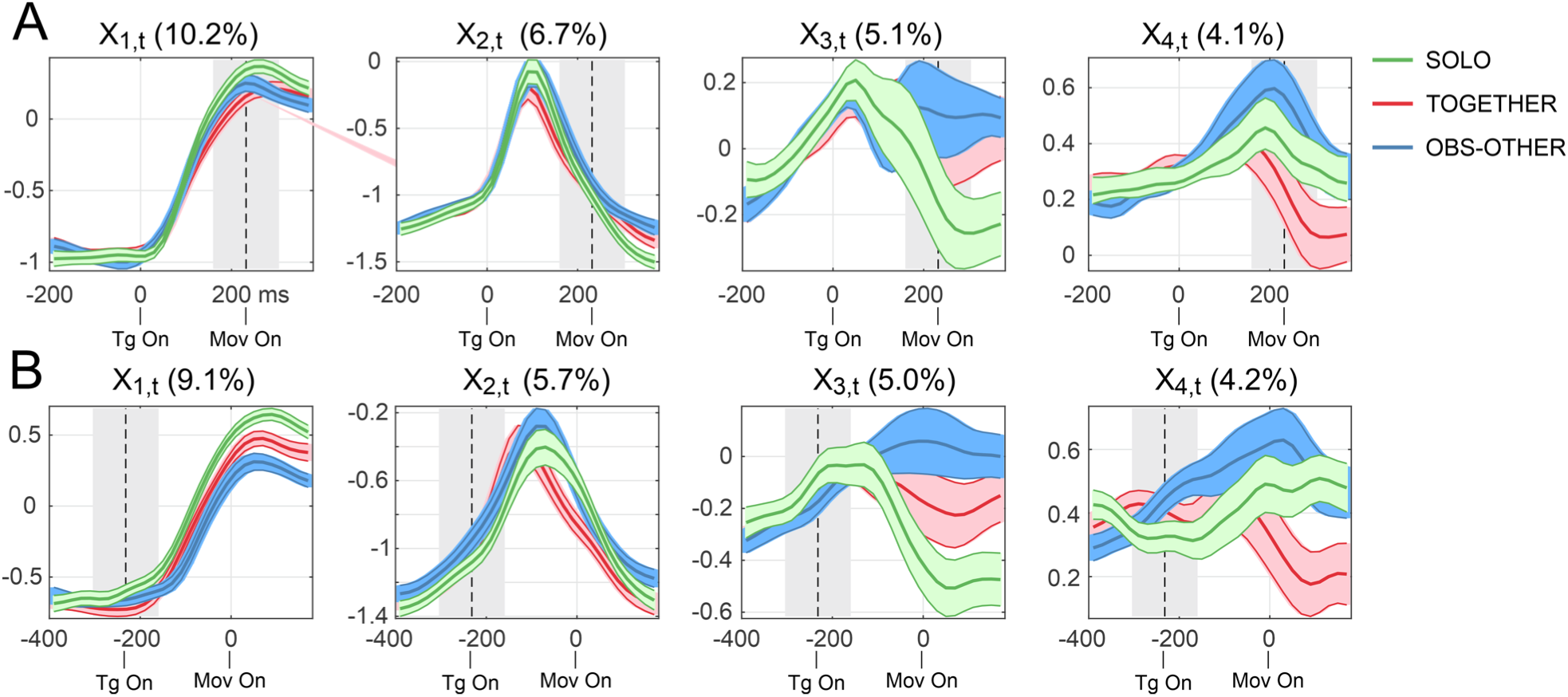
First four components of PCA aggregated according to the three task types. The four highest-ranked PCs, which captured in total 26,1% (plot A) and 34% (plot B) of the firing rate variance are plotted, after being aggregated relative to the 3 task to be performed (SOLO, OBS-OTHER and TOGETHER). The curves are obtained after aligning (0 ms) the neural activity to target onset (TgOn) (***A***) or cursor’s motion onset (Mon On) (***B***). The dotted vertical line and grey zone indicate the mean and variance (+/-SD) values of cursor’s motion onset (***A***) and target onset (***B***), respectively.

**Figure 4.**
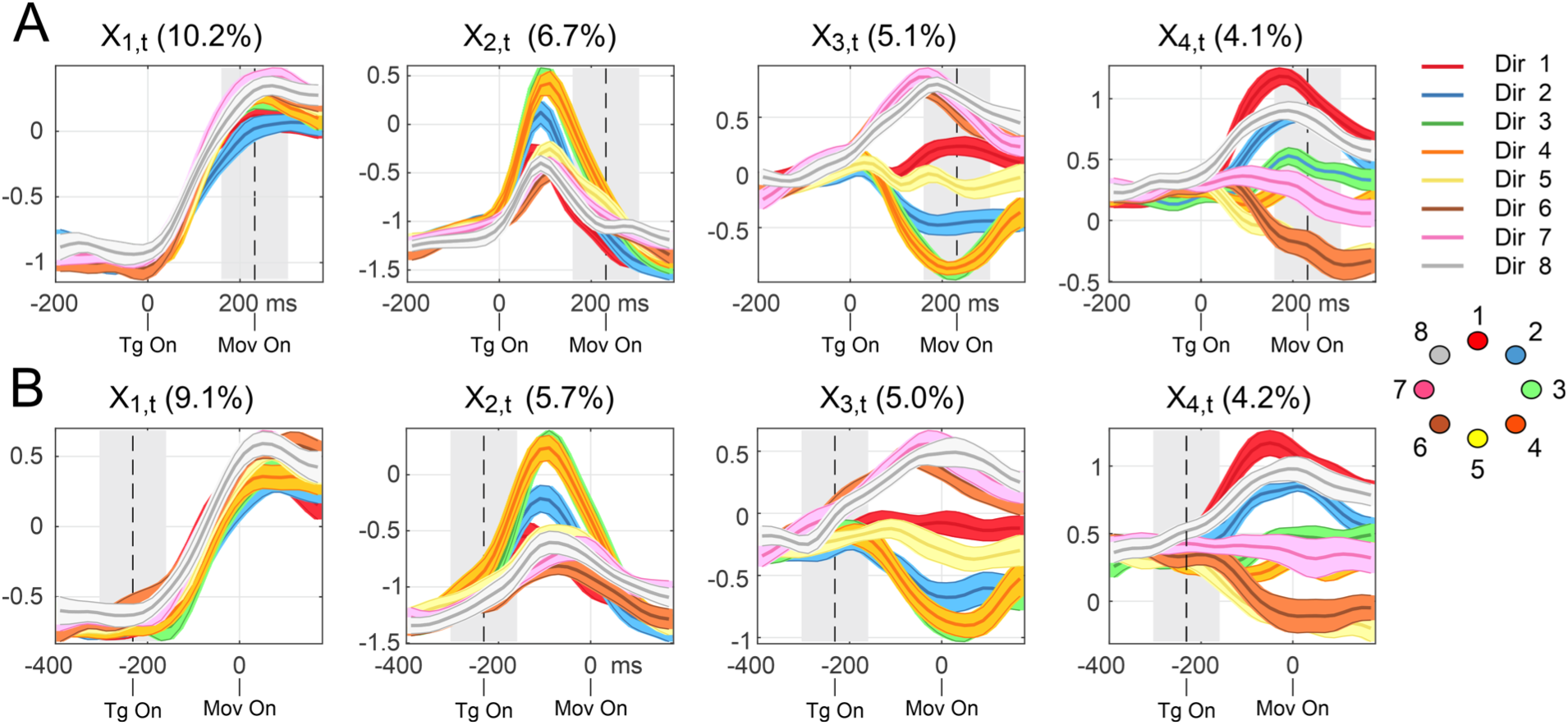
First four components of PCA aggregated according to the eight cursor’s motion directions. The four highest-ranked PCs, which captured in total 26,1% (plot A) and 34% (plot B) of the firing rate variance are plotted, after being aggregated relative to the 8 directions of cursor’s movement (Dir 1-8). The curves are obtained by aligning (0 ms) the neural activity to target onset (TgOn) (***A***) or cursor’s motion onset (Mov On) (***B***). Conventions, symbols and statistical procedures as in Fig. 3.

#### Analysis of the four highest-ranked components aggregated by task type

The profiles of the two highest-ranked principal components X1-X2 obtained by aggregating across task types (SOLO, OBS-OTHER and TOGETHER; Fig. 3) express mainly the dynamical aspects of the tasks, in a congruent manner across task types. The first component X1 shows a “state change” at target onset, whose peak is locked to the time of action initiation, occurring on average 220 ms after target presentation. The second component X2 shows a separate dynamical component, with a bell-shape profile evolving during the planning phase and peaking (irrespective of the task) at about 100 ms after target onset.

While components X1 and X2 show a similar trend between conditions, task type can be discriminated from them, particularly during the cursor’s motion. This becomes evident by considering that given the bootstrap procedure adopted to plot the graphs, the non-overlapping portions of neural space trajectories lies in the 5%-95% percentiles of the distribution of resampled differences, and consequently they can be considered significantly distinct and discriminable at p < 0.05. The distinction between tasks becomes clearer when the neural activity is aligned to MT onset (Fig. 3B). This finding indicates that while dynamical aspects are predominant, task-dependent aspects are jointly expressed with them in the two highest-ranked principal components (see also Figs. S1–S2).

The two components X3 and X4 discriminate task types even more clearly, especially around cursor’s motion onset. Notably, while components X1, X2 and X4 peak well after the target presentation, component X3 shows a different profile: its amplitude starts increasing before target presentation and peaks few tens of milliseconds after it, suggesting that it may be related to the anticipation of target onset which served as the ‘go’ signal for task initiation. Please note that the putative anticipatory effect cannot be fully explained by the time binning we adopted for data analysis, which is much shorter (20ms).

#### Analysis of the four highest-ranked components aggregated by directions of cursor’s motion

The profiles of the two highest-ranked components X1-X2 express primarily the dynamical aspects of the tasks even when aggregated across directions of cursor’s motion (D1-D8; Fig. 4). However, directional aspects can also be clearly discriminated in both. While in X1 the future direction of the cursor’s motion can be discriminated around the time of movement onset, in X2 it can be discriminated about 100ms before it, for both task alignments. Finally, the components X3 and X4 provide very effective information to discriminate all directions, even before cursor’s motion onset (see below Section 2.4).

### 3.3 Distance between trajectories in 4D neural space

#### Comparison of RMS (Root Mean Square) distances across directions and tasks

To assess the relative importance of directional and task-related information, we compared the Root Mean Square (RMS) distance in 4D neural space obtained from the first 4 components of PCA (X1-X4) of 24 trajectories (corresponding to 24 behavioral conditions: 3 task types x 8 directions), with alignment on target onset.

The Root Mean Square (RMS) distance is a standard measure of the average distance between pairs of trajectories; see the Methods for details. We performed multiple comparisons between pairs of trajectories that vary across task types and directions. For each trajectory associated to a specific task and direction (e.g. SOLO in direction D1; Fig. 5A) we calculate the distance, in any time bin, from i) the other 2 trajectories calculated in the same direction, but in the other two task conditions (i.e. SOLO in D1 vs. OBS-OTHER in D1 and TOGETHER in D1), and from ii) the other 7 trajectories calculated in the same type of task, but in different directions (i.e. SOLO in D1 vs SOLO in D2, SOLO in D3, …, SOLO in D8; Fig. 5A). The results of the 9 comparisons for SOLO in direction D1, TOGETHER condition in D1, and OBS-OTHER in D1 are plotted in Fig. 5A, 5B and 5C, respectively. The other comparisons (not shown) follow similar temporal profiles.

**Figure 5.**
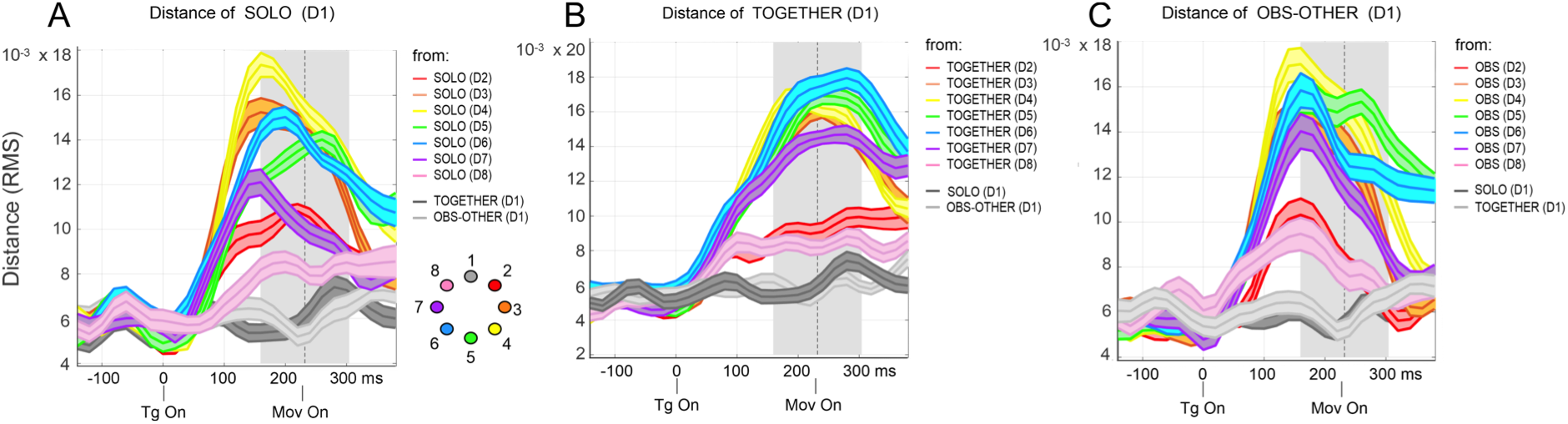
Example of comparisons of neural dynamics across tasks and directions. The comparisons are performed by computing the distance (RMS) in the 4D neural space, between pairs of neural trajetories which differ in the task (gray curves), or in the directions (colors). (***A***) RMS distances between the neural trajectory associated to SOLO condition in direction D1 vs. those differing for task but not direction (OBS-OTHER in D1, light grey, and TOGETHER in D1, dark grey) and for direction but not task (SOLO in D2, SOLO in D3, etc.). (***B-C***) The same as in A, but showing the RMD distances from TOGETHER in D1 (***B***) and OBS-OTHER in D1 (***C***). In all instances, the alignment (0 ms) of activity is to target onset (TgOn). The dotted vertical line and grey shaded area indicate the mean and variance (+/-SD) values of cursor’s motion onset, respectively.

RMS distances across both directions and tasks increase shortly (50-100ms) after target onset, but the former reach their peaks 100-150ms after target onset, well before cursor’s motion onset, whereas the latter reach their peaks after cursor’s motion onset. Importantly, neural trajectories in 4D neural space diverge more across directions of cursor’s motion than across tasks, especially before movement onset, but (for most directions) also afterwards. Furthermore, RMS distances across directions and task types show different temporal profiles, with only the former peaking before cursor’s motion onset.

The highest RMS distances are obtained when comparing the same task across directions, especially those having an absolute angular difference greater than 45° (D3-D7). Smaller differences emerge instead when comparing different types of actions, such as isometric force application vs. cursor’s motion observation (SOLO vs OBS-OTHER), or when contrasting the same actions performed in different contexts (SOLO vs TOGETHER).

To assess the generality of this finding, we repeated the same analysis for the 12D neural space, obtained from the first 12 components of PCA (X1-X12), which together captured about 40% of the firing rate variance (i.e., 41.3% and 39.6% when aligning neural activity to target onset and to cursor’s motion onset, respectively). The results of the analysis in 12D neural space are analogous to what reported for the 4D space, see Figure S7.

#### Comparison of RMS distances within motor tasks and between motor and non-motor tasks

We next assessed whether RMS distance between OBS-OTHER and the two motor tasks (SOLO and TOGETHER) is greater than RMS distance within the two motor tasks. For this, we averaged all neural trajectories belonging to the same task and calculated their RMS distances over time. Figures 6 plots the RMS distance between SOLO vs. TOGETHER, SOLO vs. OBS-OTHER and TOGETHER vs. OBS-OTHER. The figure shows that in 4D neural space, the neural trajectories begin to diverge from one another within 100 ms after target onset, in the same way across comparisons, until the onset of cursor’s motion time. At this point, the divergence amplitudes start to differ for the three comparisons.

**Figure 6.**
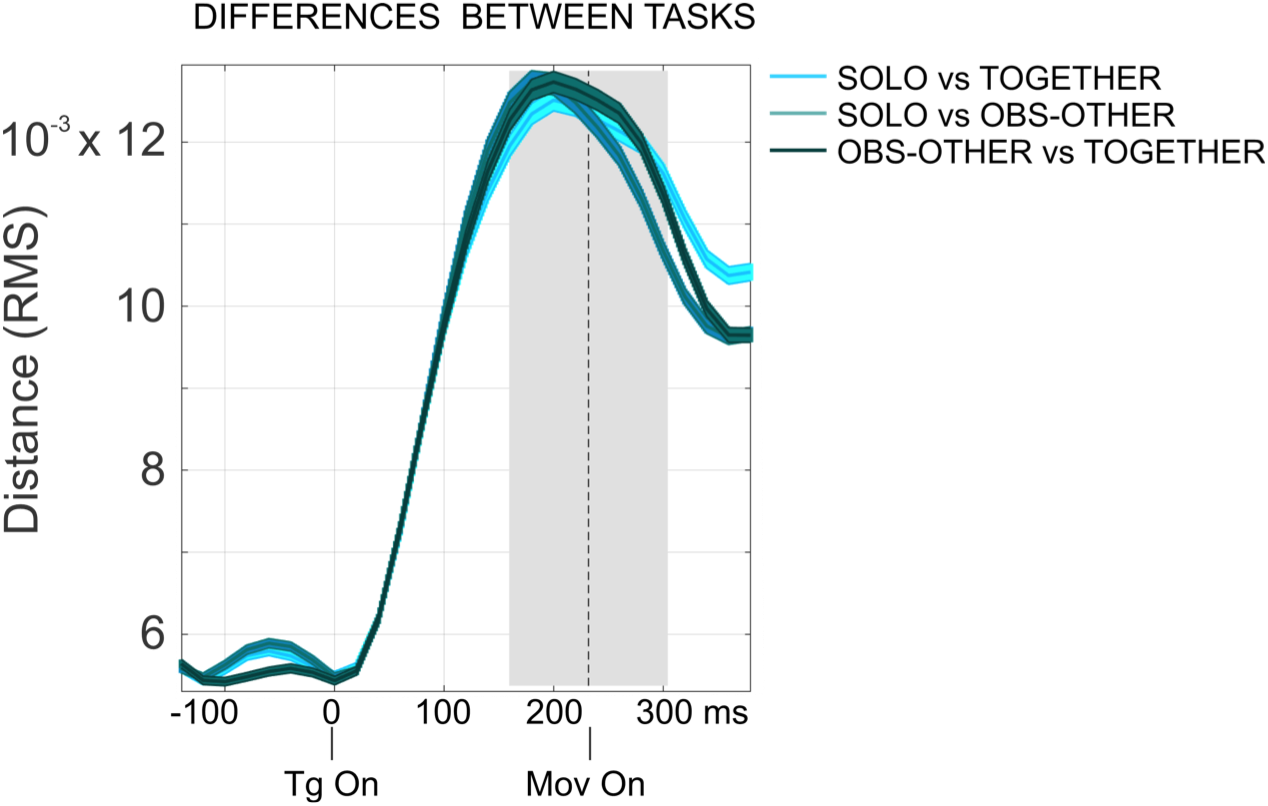
Distances (RMS) in 4D neural space between tasks. The three comparisons (SOLO vs. TOGETHER, SOLO vs. OBS-OTHER and TOGETHER vs. OBS-OTHER) are performed by firstly averaging all the neural trajectories belonging to the same task and then computing their distances (RMS) in the 4D neural space. Alignment (0 ms) is to target onset (TgOn). The dotted vertical line and grey zone indicate the mean and variance (+/-SD) values of cursor’s motion onset, respectively.

To assess the generality of this finding, we repeated the same analysis for the 12D neural space, obtained from the first 12 components of PCA (X1-X12). The analysis of 12D neural space shows a very similar trend as the 4D analysis, see Figure S8. Similar to the 4D neural space, the trajectories in the 12D neural space begin to diverge from one another within 100 ms after target onset, in the same way across comparisons, until the onset of cursor’s motion time. At this point, the divergence amplitudes start to differ for the three comparisons: SOLO diverges similarly from the other two task, whereas TOGETHER and OBS-OTHER are more different from one another than they are from SOLO.

In sum, the analysis of RMS distances in 4D neural space indicates that neural trajectories diverge earlier and to a greater extent across directions of cursor’s motion than across task types. Furthermore, and surprisingly, neural trajectories do not diverge more between motor and non-motor tasks (SOLO vs. OBS-OTHER) than within motor tasks (SOLO vs. TOGETHER).

The analysis of RMS distances across directions suggests that these distances may not be arbitrary, but rather reflect a gradient of distance in physical space between targets. For example, in Fig. 5A, the RMS distance between SOLO-D1 and its two “neighbours” in space (SOLO-D2 and SOLO-D8) remains relatively small, whereas it increases more steeply for D3-D7. Our next analysis assesses the possible isomorphism between distances across directions in neural and physical spaces.

### 3.4 Analysis of the topographical order of directional target encoding

We asked whether a topographical order exists in the representation of the directions of cursor’s motion in neural space, and whether it may reflect constraints of the external world, such as the spatial distance between the eight peripheral targets.

For each of the three tasks, we plotted the (average) neural trajectories for the 8 directions, in the 2D space formed by the two PCA components that were more direction-sensitive. Directional coding appears in all the PCs analysed in this study (Figs. S1 and S2). Hence, to evaluate which pair of components Xi, Xj (i≠j) could be considered as the best candidates to form the basis of our neural space, we calculated the pair which showed the highest performance in classifiying the cursor directions (Fig. S3). These resulted to be X3 and X4, which together explained 9.2% of the variance, when aligning the activity to both target onset (Fig. 7A) and on cursor’s motion onset (Fig. 7B). In all panels, circles, crosses and squares indicate the trajectory points corresponding to the starting, alignment and ending time adopted for plotting the curves, respectively (see Fig. 7 legend).

**Figure 7.**
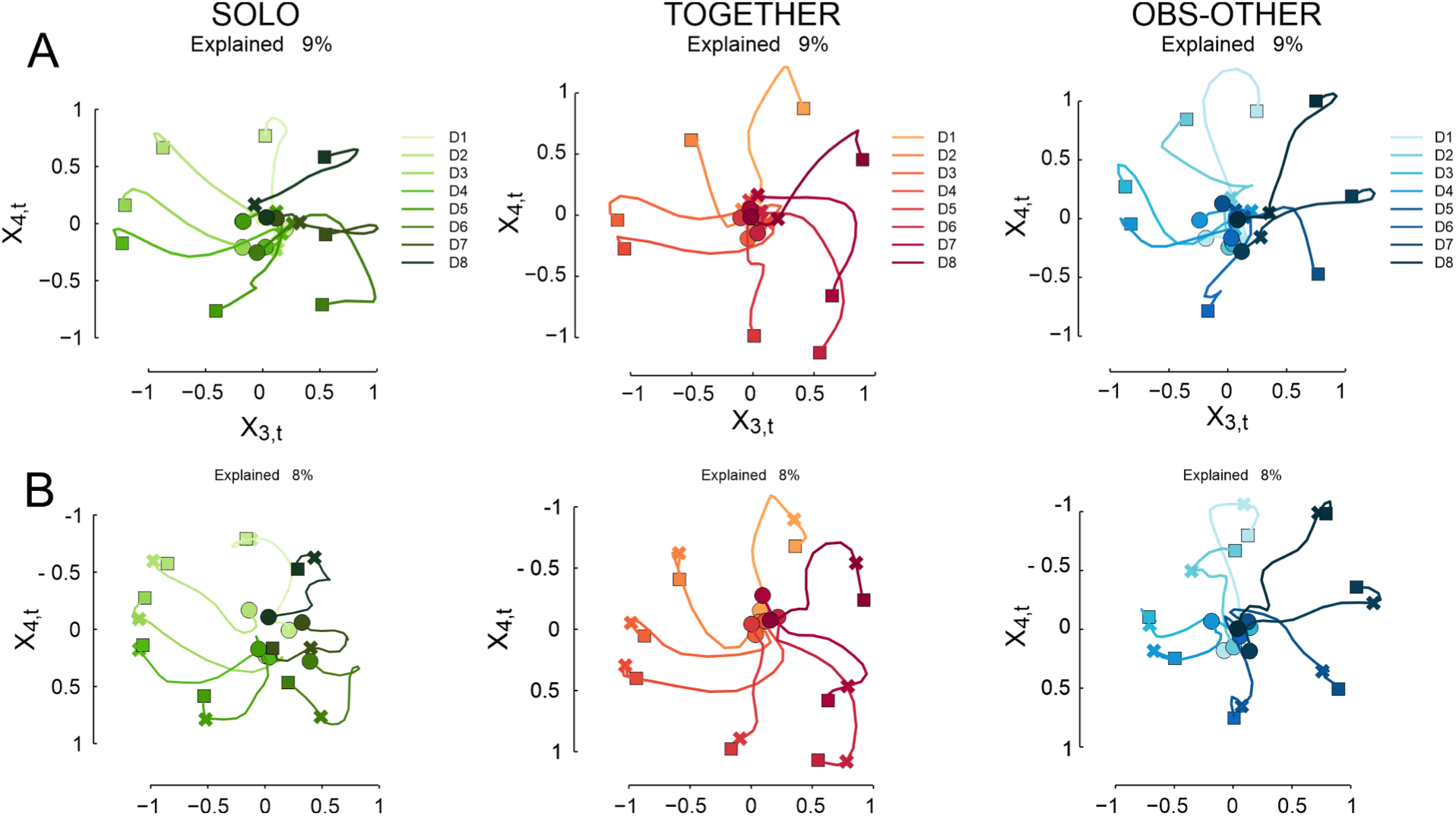
“Neural clocks”: a population-level signature of the spatial coding of target directions. Neuronal trajectories in the 2D space formed by the two most direction-sensitive PCA components: X3, X4. Each colored trajectory is obtained by averaging all the trials associated to a given target direction (D1-D8), in the SOLO (***A,B***; green), TOGETHER (***A,B***; red) and OBS-OTHER (***A,B***; blue) tasks. (***A***) The neural trajectories are aligned to target onset and the interval shown is [-200ms, 400ms]. Circles mark the starting point, 200 ms before target onset; cross symbols indicate target onset (0 ms) and squares mark the end of the interval, 400 ms after target onset. (***B***) The neural trajectories are aligned to cursor’s motion onset and the interval shown is [-500ms, 200ms]. Circles mark the starting point, 500 ms before target onset; cross symbols indicate cursor’s motion onset (0 ms) and squares mark the end of the interval, 200 ms following the cursor’s motion onset. Note that each point in the neural trajectory is the average of 20ms of neuronal activity, e.g., point 0 is effectively the interval [-10ms,10ms].

In all the three tasks, the neural trajectories are clustered together in a common space before target presentation (circles; Fig. 7A), as previously reported in studies using a delay period ^21^. All the trajectories diverge already around target onset (crosses; Fig. 7A), well before cursor’s movement onset (crosses; Fig. 7B). Interestingly, the neural representation of cursor’s motion in the 8 directions in neural space has a topographical structure. Neural trajectories are organized according to a directional gradient that is isomorphic to the spatial arrangement of target positions in the external world. In other words, targets that are closer in external spatial coordinates (e.g., D1 and D2 but not D6) are closer in neural space, too. Here, we refer to this directional gradient as a “neural clock” to highlight the resemblance with an analog clock face (please note that in this context, the term “clock” has nothing to do with temporal representations).

To quantify the quality of the directional gradients, we calculated a measure of angular distance between the 8 neural trajectories aligned to target onset, by considering the eight lines that start from the origin (e.g. the center of Figure 7A in the SOLO task) and arrive at the end of each neural trajectory (e.g., the dark green square in Figure 7A that marks the end of D8 in the SOLO task). Then we checked if, for each neural trajectory (e.g., D8) the smaller angular distance in the clockwise and anticlockwise directions are with the “neighbour” neural trajectories (e.g., D7 and D1, respectively) or not. This results in 16 comparisons for each task: 8 neural trajectories (D1-D8) by 2 neighbours (clockwise and anticlockwise). The test of the comparisons resulted in a perfect score (16/16 successes) for all the three tasks, indicating that the directional gradients are topographically ordered.

Intriguingly, the same (counter-clockwise) directional gradient is present not only in the two motor tasks, but also during action observation, when monkeys do not control the cursor’s motion or apply any hand force. A control analysis of activity during a “saccade only” task (see Figs. S4 and S5) rules out the possibility that this similarity is due to oculomotor behavior, which has been reported to be similar across all the task conditions ^26,35^.

## 4. DISCUSSION

We studied population-level dynamics in the premotor cortex of monkeys that performed two isometric tasks and an observation task. During the isometric tasks, monkeys applied a hand force on an isometric joystick to guide a visual cursor from a central position toward one of eight visual targets, individually (SOLO) or together with another monkey (TOGETHER). During the observation task, each animal observed the moving cursor controlled by its partner (OBS-OTHER), when the latter perfomed its SOLO trials.

We used a dimensionality reduction (PCA) analysis to ask (i) how task-related variables (e.g., action initiation, cursor’s motion direction, task type) are encoded at the population level and (ii) whether this population coding differs when contrasting different behavioral demands, such as the two motor tasks (SOLO and TOGETHER) versus the non-motor task (OBS-OTHER).

### 4.1 High-ranked components signal shared dynamics across behavioural tasks

The multi-task approach of this study afforded different comparisons of the neural dynamics across several cognitive-motor conditions. In particular, we contrasted: i) similar motor tasks performed in two different contexts (e.g. SOLO vs TOGETHER); ii) action execution and observation (e.g. SOLO vs OBS-OTHER); iii) same type of action performed in different directions (e.g. SOLO in eight directions); and iv) observing the consequences of the other partner’s action, i.e. a visual cursor’s motion, toward different locations (i.e. OBS-OTHER).

The analysis of the highest-ranked principal components that emerged from PCA (i.e., those explaining the most variance) showed that in premotor cortex a large portion of the neural variance was shared (i.e., had similar dynamics) across the three tasks. This is despite the fact that the three tasks involved significant differences in behavioral demands. Motor (SOLO and TOGETHER) and non-motor (OBS-OTHER) tasks require the execution or only the observation of movement, respectively. The two motor tasks (SOLO and TOGETHER) require exerting the same amount of force on the joystick to reach the target, but provide slightly different visual inputs during the performance (e.g., one cursor vs. two cursors with a circle). They also have different control demands and attentional load, with the TOGETHER tasks requiring accurate coordination of movements with the partner.

Previous studies of neural population dynamics in motor and premotor cortex that used dimensionality reduction (like PCA) also found the highest ranked components to be related to dynamical aspects of the task; and interpreted them as related to state changes and movement timing, rather than movement type ^36^. Views of motor cortical activity operations regarded as a dynamical system have proposed that a major proportion of premotor activity is devoted to signaling phase dynamics and in particular preparatory states, reflecting the initialization of a dynamical system at optimal conditions, which will later generate a movement ^21,22,37^. Our findings confirm the predominance of dynamical aspects in the two highest-ranked components of PCA (X1 and X2), but offer a new neurophysiological interpretation compared to those proposed so far in the literature.

#### Dynamical aspects of neural space do not necessarily reflect an internal translation from movement preparation to its generation

The similarity of population-level coding across tasks that require and do not require movement may help in contextualizing the interpretation of previous findings about action coding in premotor cortex, and especially the roles of the dynamical aspects of neural space found to be encoded in the highest-ranked PCA components. Our findings are in fact consistent with previous works using a range of techniques, including the orthonormalized neural trajectories of a Gaussian Process Factor Analysis shown in Yu et al. ^32^, the Canonical Correlation Analysis (CCA) projections of the neural population response and the modeled data ^38^ and the dPCA components of Kaufman et al. ^36^. As a reminder, the CCA attempts to find the patterns which are common across two data sets, such that the reweighted data sets (i.e. canonical variables) are maximally correlated.

In Yu et al. ^32^ and following studies ^36,39^, the interpretation of dynamical aspects of neural space is that their pattern is “presumably related to generating the arm movement and it is thus sensible that it is time-locked to movement onset”. However, in the above studies, the results were obtained only from neural data collected during motor performance in a classical delayed-reaching paradigm. As these studies did not investigate the dynamics of forms of behavior not requiring movement generation, the proposed interpretation might have been biased by the nature of the adopted experimental paradigm.

Here, instead, we considered also a condition requiring monkeys to observe the consequences of the motor output of another monkey. Our findings indicate that in premotor cortex the same dynamics found in two isometric tasks (SOLO and TOGETHER) is shared with a behavioral condition which did not involve a motor act, but rather its abstract representation (OBS-OTHER). We can rule out the possibility that the similarity between motor and non-motor tasks is due to subtle movements that the animal may have performed during the observation task, because no such movements emerge from the analysis of behavioral data (see the speed profile of the cursor, purple curves in the bottom panels of Fig. 2). Furthermore, we can exclude that the similarity between motor and non-motor tasks can be attributed to eye behavior. To this purpose, we performed a control PCA analysis that considers the three main tasks (SOLO, TOGETHER, OBS-OTHER) along with a fourth task (EYE task) that involved the same oculomotor behavior, but no action execution or cursor motion observation; see ^26,35^ for more details about the similarity of eye movemens across tasks. The analysis of the highest-ranked principal components X1-X4 of the control PCA on the four tasks reveals that the neural representation of the EYE task is markedly different from the other three ones (see Figs. S4 and S5). To our knowledge, this is the first investigation of the putative influence of oculomotor behavior on the neural dynamics associated with motor behavior.

The fact that action observation leads to a pattern of highest-ranked components X1 and X2 similar to those associated to action requiring hand force application offers a different perspective on their roles – which goes beyond their reiterated attribution to movement generation ^32,36,39^, which was also predicted by theoretical models ^38^. Kaufman et al. ^36^ suggested that highest-ranked components reflect an internal transition from movement preparation to movement generation. Our data in the OBS-OTHER condition suggest that the transition does not necessarily imply a physically performed movement, but also an internally simulated action.

Alternatively, one may consider the temporal profile of highest-ranked components found in our and other studies to correspond to “condition-independent” dynamics – a view consistently proposed to interpret the results of previous studies in different cortical areas and different tasks ^36,39–41^. Condition-independent components have been regarded ^40^ as the building blocks of neural activity, which capture its temporal modulations throughout the trial; and similar to our study, they are mostly locked to stimulus presentation. In this study, it is plausible that these components replicate the analogous sensory-motor temporal structure shared in the three behavioral conditions (SOLO, TOGETHER, OBS-OTHER). The dynamics captured by the pattern of the two highest-ranked components might therefore mirror the occurrence or the expectancy of behavioral events (e.g. central target appearance, followed by a peripheral target presentation, which in all instances predicts a cursor’s motion), or more simply the presentation of a task-relevant stimulus – which are shared across tasks. Finally, we cannot exclude that the condition-invariance of the highest-ranked components might relate to decision dynamics occurring between the peripheral target presentation and cursor’s motion onset, which select the type of action and the direction of the (real or internally simulated) force application. In this perspective, the neural dynamics extracted from the higher-order components may reflect evidence accumulation ^42^ or an urgency signal ^43,44^ that guides the choice between different action types, rather than motor generation. These and other alternative hypotheses remain to be fully explored in future studies.

### 4.2 Directionality encoding

We found that that direction of cursor’s motion and task identity can be decoded prior to cursor’s motion onset in the highest-ranked components and intertwined with phase dynamics (Fig. 3–4), rather than being only expressed in lower-ranked components. Our PCA-based analysis hence recovers the results of numerous previous studies that showed that spatial and motor parameters such as movement direction ^45,46^ and force can be accurately decoded from cortical populations ^47–49^.

Interestingly, neural trajectories diverged more profoundly across directions of cursor’s motion than across task types for any given direction (see Fig. 5). In other words, in “neural space”, it is easier to discriminate trajectories that vary across different directions (e.g., D1 in the SOLO task versus D2-D8 in the SOLO task) than across task types (e.g., D1 in the SOLO task versus D1 in the TOGETHER or OBS-OTHER tasks). This result suggests that spatial coding may consist of a neural process that operates separately and independently from a neural code aimed at distinguishing different types of action. This separation might not be expressed at the level of single components, where spatial and task-type information can be mixed, but rather emerges in the analysis of 4D neural space.

A rationale for this organization of neural space may lie in the functional specialization of premotor cortex, in which dynamical and spatial aspects of movement may need to be expressed independently from the specific task to be performed. It remains to be investigated in future research whether dynamical, spatial and task-related aspects are expressed by the same population of premotor neurons or by different subpopulations. In this latter case, it is possible to speculate that one subpopulation of neurons may convey dynamical and kinematic features to shape motor representations that can subserve different contexts and tasks, including action planning, execution and observation; whereas other subpopulations of neurons (whose variance is captured by low ranked PCA components) may convey higher-order signals for task- and context-related information, which allow distinguishing amongst behaviors with similar temporal structure, like SOLO, TOGETHER and OBS-OTHER. This latter scenario would be in line with a *single neuron network topology*, based on the existence of functional modules across cortical areas that guarantee fast and dynamical information processing and transfer ^50^, and where the operations of even a small fraction of the network are sufficient to characterize a reasonable amount of the spatio-temporal features of neural activity.

#### Neural clocks: isomorphism between target direction representation in neural and external spaces, across motor and non-motor tasks

We found a remarkable isomorphism between the neuronal representation of movement directions and the arrangement of targets in the external space (see ^51^ for a related finding). Similar to previous studies, the trajectories in the 2D neural space of Fig. 7 start from a central, equi-potential (or non-informative) state and diverge at the target onset ^21,52^. However, remarkably, they follow directions in neural space that are isomorphic to the spatial position of external targets. This topographical organization (“neural clock”) may be a potential population-level signature of the spatial coding of target directions. Interestingly, the “neural clock” emerges in all the three tasks; and has the same orientation with respect to the real physical space of targets. In contrast, it does not emerge in a control task consisting of saccadic eye movements to eight spatial targets, suggesting that it does not result from oculomotor dynamics (see Figs. S4 and S5).

## 5. Conclusions

In sum, the present results indicate a novel sharing of population-level dynamics in monkey premotor cortex across tasks that required movement generation (motor tasks) and those that did not require it (non-motor tasks), despite the strikingly different behavioral demands. The sharing becomes evident by considering the similarity of the highest-ranked components and of the “neural clocks” across the three tasks considered in this study; and the fact that trajectories in neural space do not diverge more between motor and non-motor tasks than across motor tasks. These findings suggest that the largest components of population dynamics in premotor cortex express dynamical and spatial aspects of movement independently from the task to be performed and hence they are not necessarily implicated in the translation from movement preparation to its generation. The similarities we found between population dynamics in PMd across tasks that required and did not require movement generation suggests that they express the temporal and directional features of moving the cursor to the goal rather than any actual movement per se. Therefore, this study contributes to a large body of evidence showing cognitive roles of population dynamics in premotor cortex (e.g., motor imagery, decision-making, and discrimination ^7,11,53–55^), by suggesting that the highest ranked components of premotor population activity may reflect an abstract, covert representation of action and its goal rather than mechanisms strictly confined to motor functions. The involvement of the highest ranked components in cognitive functions suggests that the latter are central to premotor population dynamics rather than ancillary or subordinate to motor roles.

The sharing of action representations across motor and non-motor task reported in this study is consistent with a large body of literature. First, single cell data analysis of our database showed similarities of the neural activity recorded during visual observation and isometric hand action trials^26^, with one third of the original dataset consisting of cells modulated during both observation and at least one of the two isometric action tasks. Despite the heterogenity of neural responses, quantified on the basis of their firing rates, some cells were similarly modulated or shared directional tuning across task conditions. Furthermore, our findings are reminiscent of mirror-like mechanisms, possibly supporting a matching operation between action execution and observation ^14,15,56,57^ and are in line with other recent studies indicating that in motor and premotor cortex, population dynamics share a common neural subspace between action execution and observation ^20^. We note, however, that because we used isometric tasks, what monkeys actually observed are the *consequences* of the companion’s actions, not their overt movements. The putative matching between performed and observed actions occurs at the level of the cursor’s motion, not of limb movements - similar to the shared representation found in premotor neurons in abstract contexts akin to ours ^16,17^. Finally, our results are in keeping with the finding that dorsal premotor cortex preferentially codes target position and not limb movement during a delay period ^58^; see also ^59^. Our study suggests that a similar mechanism could operate during reaction-time tasks, without explicit delay periods. Previous evidence of similarities across motor and non-motor conditions were interpreted in terms of movement suppression ^60^ or action simulation ^11^. The present study was not designed to disentangle between these (or other) possibilities, which require further investigation.

## Acknowledgements

This research received funding from the MIUR of Italy, Prin 2017 (Grant N. 201794KEER_002) to ABM; the SAPIENZA-University of Rome (“Ricerche universitarie 2019”) to ABM; the European Union’s Horizon 2020 Framework Programme for Research and Innovation under the Specific Grant Agreement Nos. 785907 and 945539 (Human Brain Project SGA2 and SGA3) to GP; and the European Research Council under the Grant Agreement No. 820213 (ThinkAhead) to GP. The authors are sincerely grateful to Prof. Roberto Caminiti for his valuable suggestions and remarks.

## SUPPLEMENTARY MATERIALS

### Classification of task types and directions from principal components

We conducted a further analysis to corroborate the finding that information about task types and direction is present across multiple principal components, rather than localized only in some of them (e.g., those ranked as the highest). To this aim, we computed the probability P(C|X) to correctly classify the class C given a PC-specific dataset of trials X. We made 12 PC-specific datasets, one for each PC component X1-X12. Each trial in the database X is a is a sequence of PC-specific coefficients c(t0), …, c(t_end) in the same temporal interval [t0,t_end] considered in the main analysis. In Figure S5, the classes C are the three task types (SOLO, TOGETHER, OBS-OTHER). In Figure S6, the classes C are the eight directions D1, …, D8. For each component, we treated each of the 192 trials as a vector having one label for task type (“together”, “solo” or “obs-other”) and one label for direction (“1” to “8”). We used naïve Bayes to compute the probability to classify each trial correctly for task type (chance level is 0.333) and direction (chance level is 0.125) separately.

Figure S1 plots the average and confidence interval of the probability of correctly classifying trial types, for each component X1-X12. The figure shows that all tasks can be classified above chance (horizontal dotted line) in almost all the principal components, including the highest ranked components, albeit with some differences (e.g., X1 classifies “together” and “solo” better than “obs-other”; X2 classifies “obs-other” and “solo” better than “together”).

**Figure S1.**
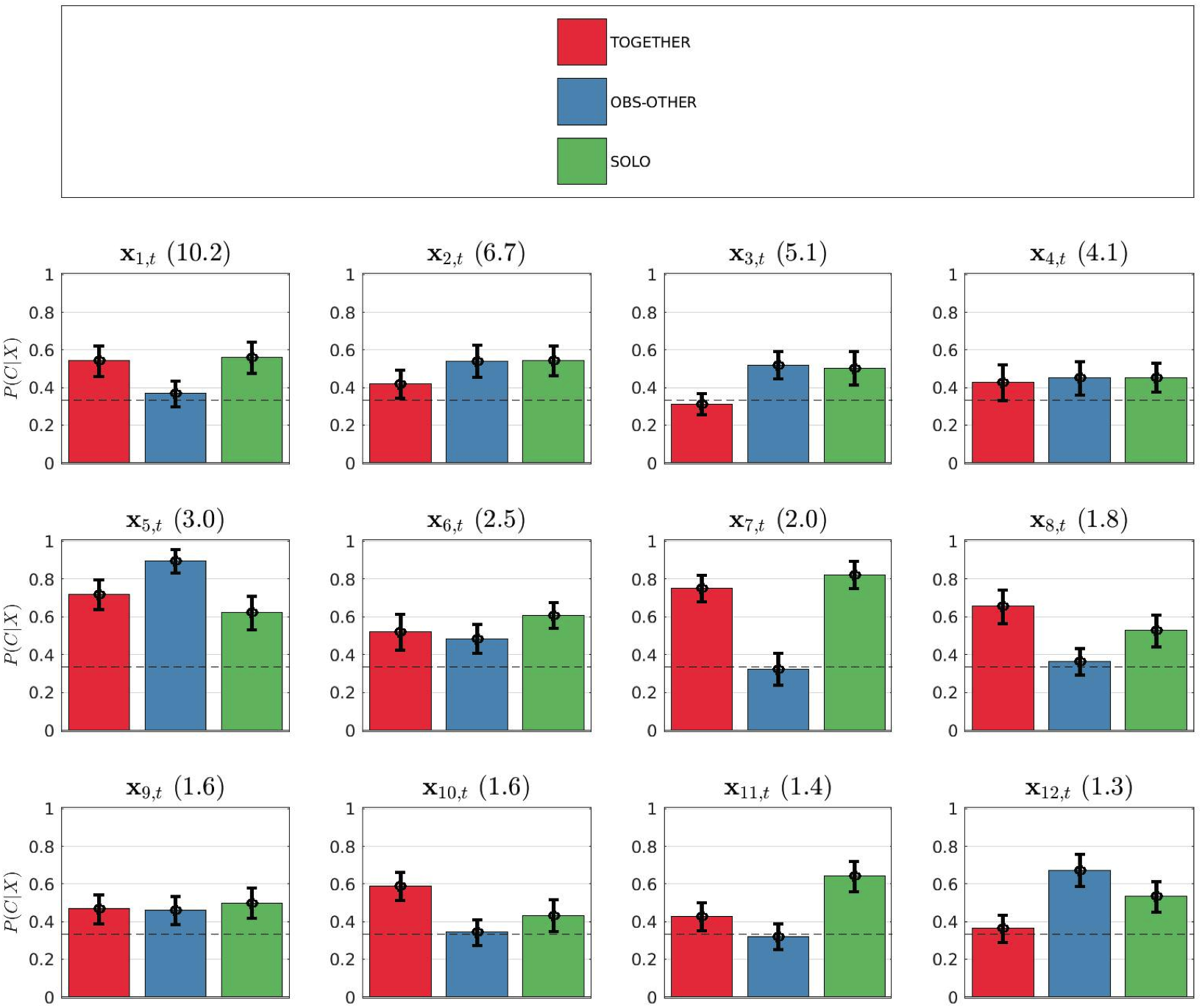
Probability of the correct classification of task types for each principal component X1-X12. Each colored bar shows the average and confidence interval of P(C|X) for one task type (“together”, “solo” or “obs-other”). The horizontal dotted line indicates chance level (0.333).

Figure S2 plots the average and confidence interval of the probability of correctly classifying directions, for each component X1-X12. The figure shows that all directions can be classified above chance (horizontal dotted line) in almost all the principal components, including the highest ranked components, albeit with some differences (e.g., X1 classifies directions 1 and 2 better than the other directions; X2 classifies all directions, but direction 7 with lower probability).

**Figure S2.**
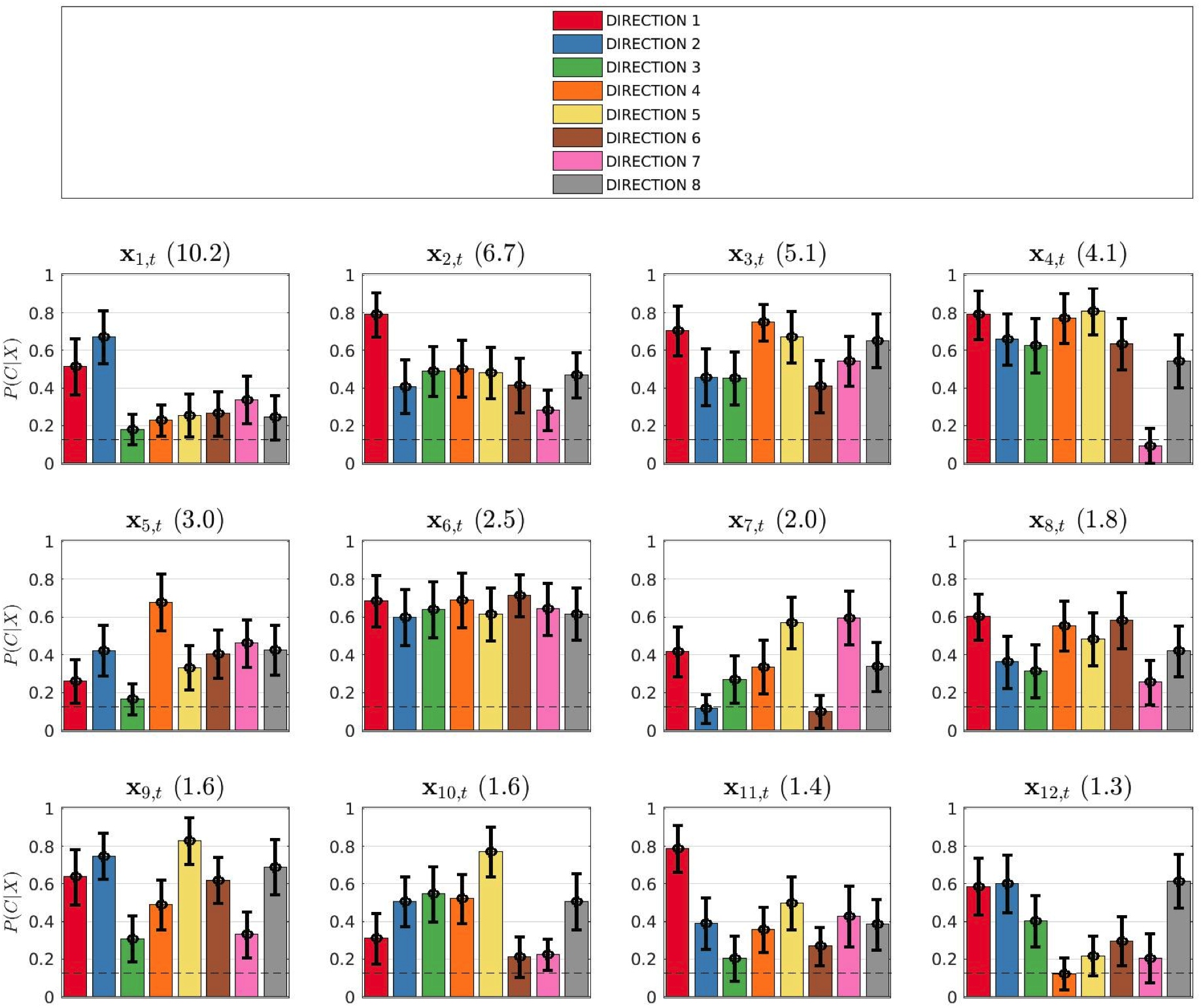
Probability of the correct classification of directions for each principal component X1-X12. Each colored bar shows the average and 90% confidence interval of P(C|X) for one direction (1 to 8). The horizontal dotted line indicates chance level (0.125).

### Control analysis to determine which pair of PCs better classify directions

To plot the “neural clock” shown in Figure 7, we analysed which pair of components Xi, Xj (i≠j) - taken together - where the most direction-sensitive and hence optimal candidates to form the basis for computing the 2D neural trajectories. To this aim, we calculated which pair of principal components better classified the directions. We used the same classification methods as for Figs. S1 and S2, but considered the coefficients of pairs of PC components rather than single PC components.

Figure S3 plots the result of this analysis, limited to the highest components X1-X4, to maintain a high level of explained variance. The figure shows the average and confidence interval of the probability of correctly classifying directions, for all the six pairs that we considered (“X1 and X2”, “X1 and X3”, etc.). The colored bars show the probabilities of correct classification for each direction separately, whereas the black bar shows the average across directions for each pair. We found that the pair of components that classifies better the directions is “X3, X4”, for which we obtained a classification performance of 87%, with 90% confidence interval CI (83,91).

**Figure S3.**
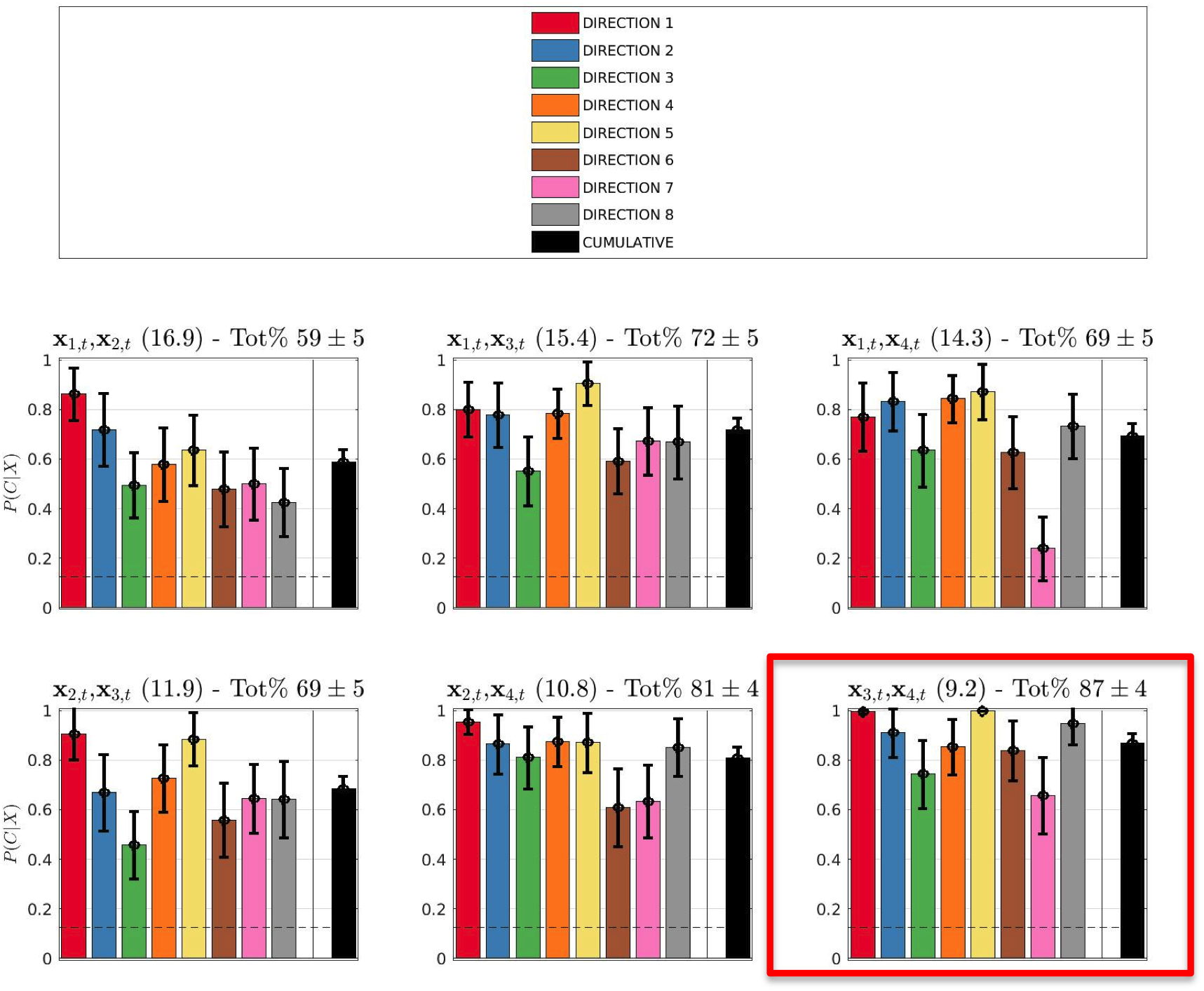
Probability of the correct classification of directions for each pair of components “X1 and X2”, “X1 and X3”, etc. Each colored bar shows the average and confidence interval of P(C|X) for one direction (1 to 8). The black bars show the grand average (and confidence interval) across all directions. The horizontal dotted line indicates chance level (0.125).

We performed an additional analysis to rule out the possibility that the information that the components “X3, X4” carry about directionality is confounded by the fact that cursor velocities may vary in the 8 different directions during task execution. For this, we performed a correlation between the peak velocities of the cursor in the 8 directions (averaged across tasks) and the probabilities to correctly classify directions reported in Figure S2, for the two components X3 and X4. We found the correlations to be very small for both X3 (R=0.032) and X4 (R=0.011), permitting us to rule out the possibility that cursor velocity confounds the directionality information of these components.

### Control analysis: eye movement task

We performed a control analysis to rule out the possibility that the common pattern of directional gradient observed in the three tasks was mainly related to eye behavior, which was virtually identical in the SOLO, TOGETHER and OBS-OTHER tasks ^2,3^. For this, we conducted a separate PCA analysis that considered data from the three aforementioned tasks along with a fourth, control saccadic task (EYE). In the EYE task, monkeys had to perform saccadic movements from the central target to one of the 8 visual stimuli presented in same locations as those used in the SOLO, TOGETHER and OBS-OTHER. Neural activity in the EYE task was recorded from the same population as the other three tasks; but while the latter were performed interminging the different types of trials in one block, the saccadic trials were presented in a separate block, before or after the performace of the block of SOLO, TOGETHER and OBS-OTHER trials. For this new PCA, the neural activity was aligned to target onset.

Figure S4A shows the four highest-ranked principal components X1-X4 obtained by clustering across task conditions (SOLO, OBS-OTHER, TOGETHER, and EYE). It shows that the neural representation of the EYE task is markedly different from the other three tasks, despite their similar oculomotor demands. The first principal component of PCA, which explains almost one third of the variance (X1, 30.7 %), clearly distinguishes EYE from the other three tasks, even before target onset; whereas the other three tasks are closely grouped together. This component may therefore discriminate only the tasks conditions performed in separate blocks. However, the other high-ranked components X2-X4 also clearly discriminate the saccadic (EYE) trials from the others, which show similar temporal trends. Figure S4B shows the four highest-ranked principal components X1-X4 obtained by clustering across directions (D1-D8). In the first principal component, the spatial features of the tasks are smeared out.

**Figure S4.**
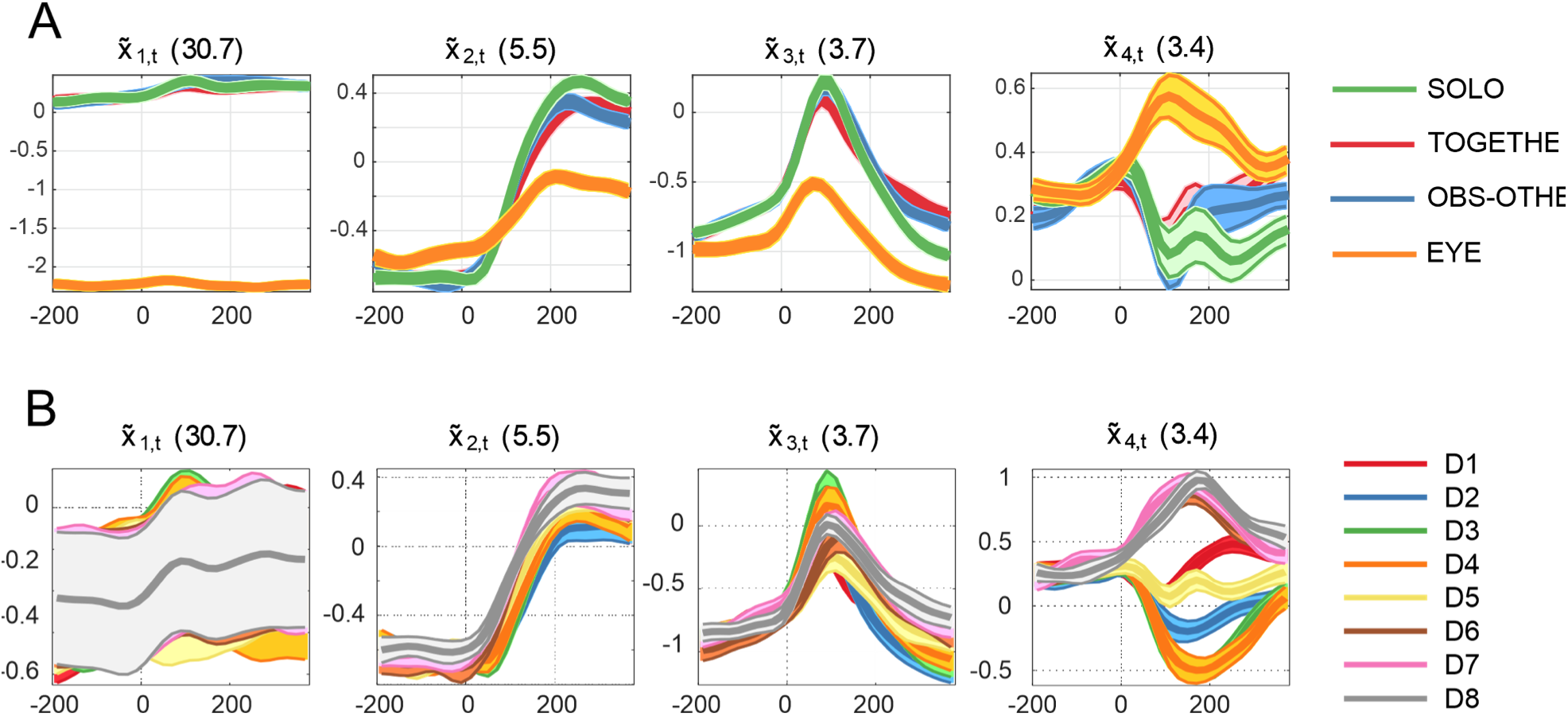
Control PCA, including eye-related activity. The first four principal components aggregated by tasks (A) and directions (B) are plotted when including neural activity studied during a control EYE task, consisting in saccadic movement in 8 directions.

Finally, although single unit activity recorded during the saccadic task was directionally tuned ^2,3^, in PCA space the activity during the EYE task did not produce the systematic directional gradient (Figure S5D) that was clearly observed in the other tasks (Figure S5A-C). To quantify the quality of the directional gradients, we used the same method based on angular distances reported in the main paper. The test of the comparisons resulted in a perfect score (16/16 successes) for the SOLO, TOGETHER and OBS-OTHER tasks but in a poor, near-chance score (4/16 successes) for the EYE task.

**Figure S5.**
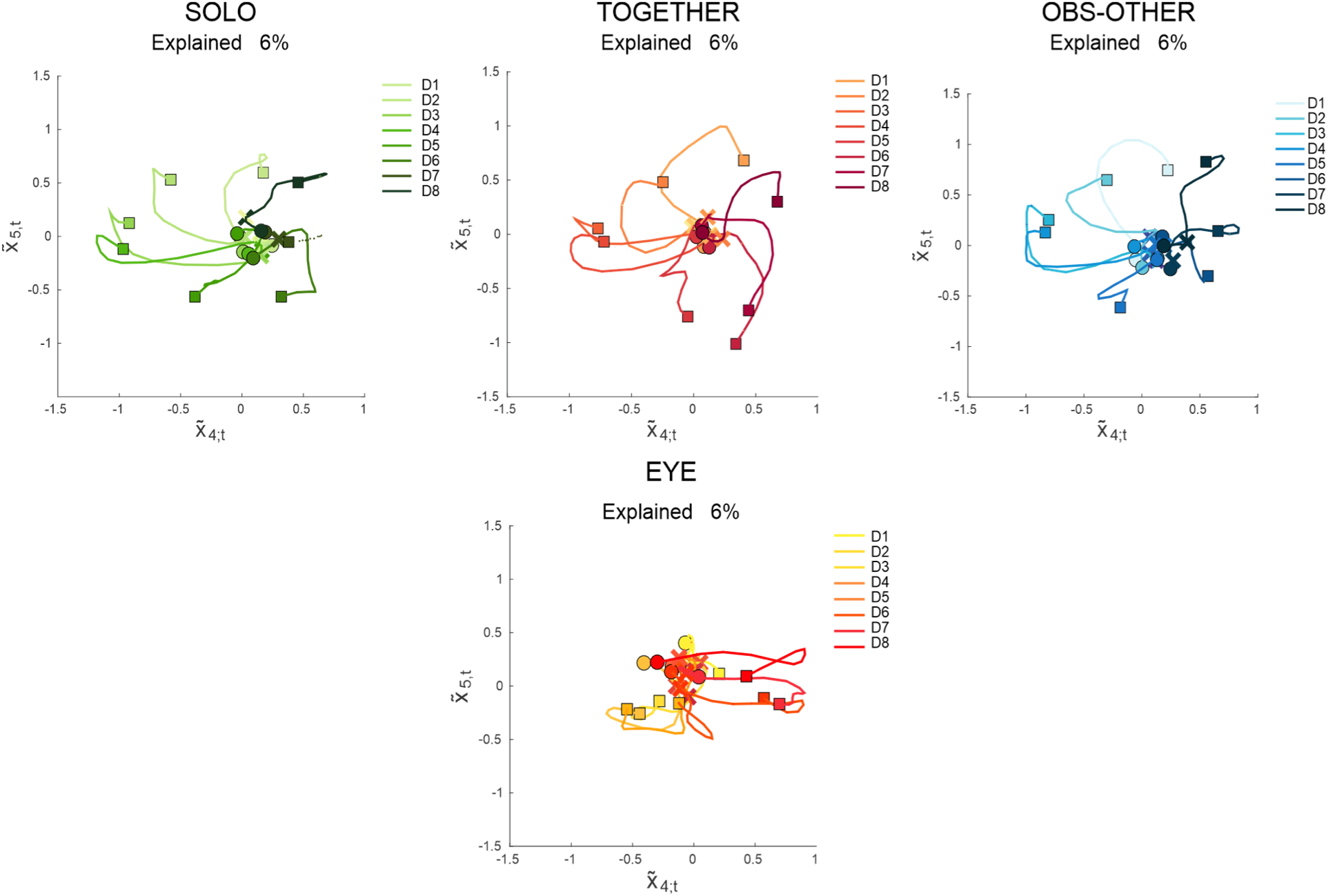
Control PCA, 2D neural trajectories obtained combining components X4 and X5. Note that X4 and X5 in this analysis are analogous to X3 and X4 in Figure 7, since X1 is a novel PC that distinguishes task blocks, see Figure S4.

### Example data from single monkeys

In the main analyses, we constructed a neural population using data pooled from 2 monkeys and 36 sessions. The pooling procedure from two animals was motivated by the fact that they were tested together, either when acting jointly, or when one acted and the other observed, and also because when individual data sets were analysed, similar results were obtained between the two. We observed minor differences in the rank and coding of the principal components. Examples of single-monkey analyses are reported in Fig. S6. It is noting the similarity of neural dynamics. While the sign of PC X2 is inverted in monkey K, the sign is arbitrary in PCA and only the relative magnitudes are meaningful ^1^ when studying the overall patterns of neural dynamics.

**Figure S6.**
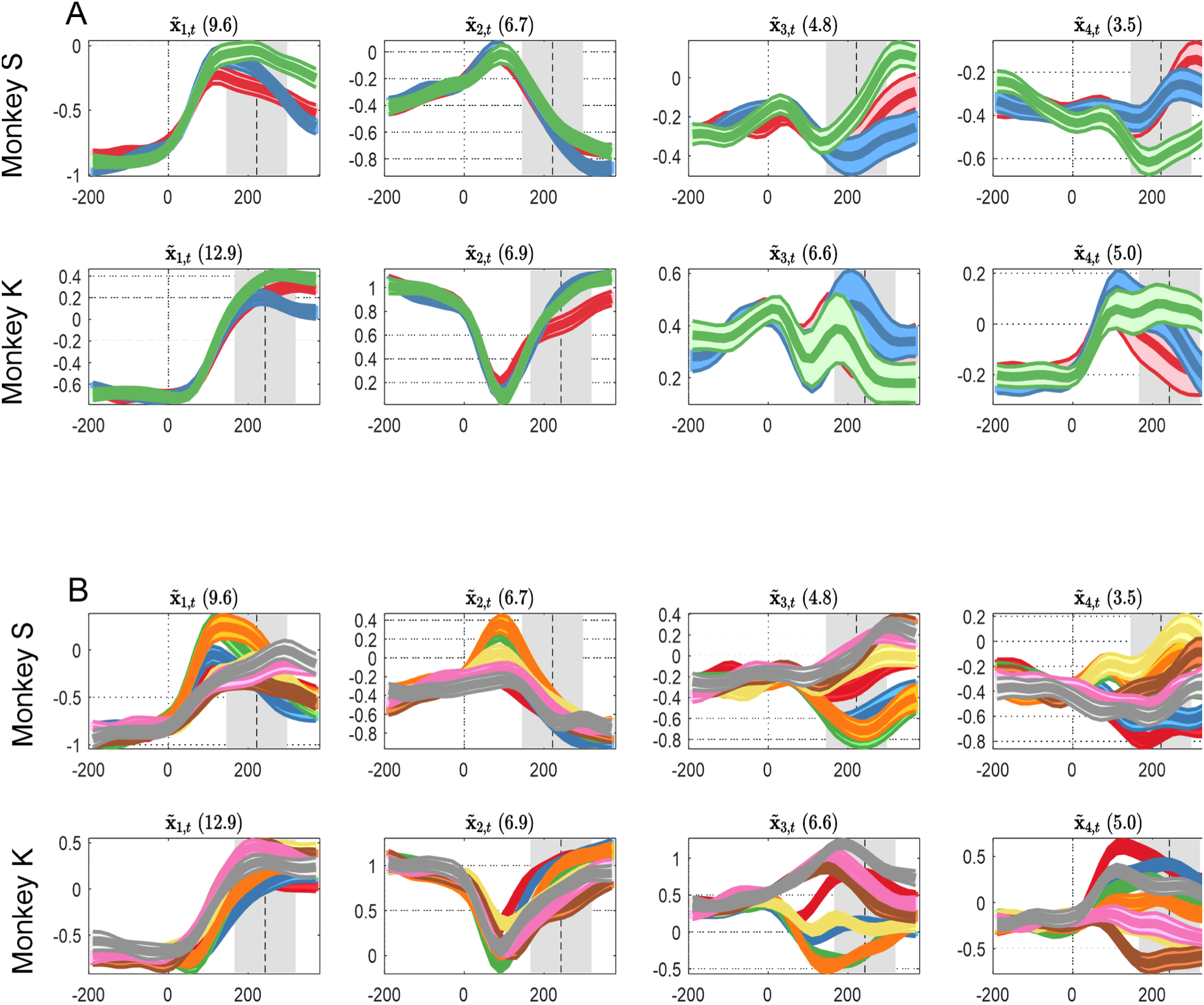
PCA obtained for each monkey. The four highest ranked PCs, aggregated by tasks (A) and directions (B), are reported for each monkey dataset analysed separately (monkey S, top panels; monkey K, bottom panels), after aligning the neural activity to target onset (0 ms).

### Distance between trajectories in 12D neural space

We conducted a control analysis of the distance between trajectories in 12D neural space, which is analogous to the analysis of 4D neural space reported in the main text, except that we considered the Root Mean Square (RMS) distance in 12D neural space obtained from the first 12 components of PCA (X1-X12) of 24 trajectories (corresponding to 24 behavioral conditions: 3 task types x 8 directions), with alignment on target onset.

The results of the comparisons in 12D neural space are reported in Figures S7 and S8, which are the equivalents of Figures 5 and 6 in the main text, respectively.

**Figure S7.**
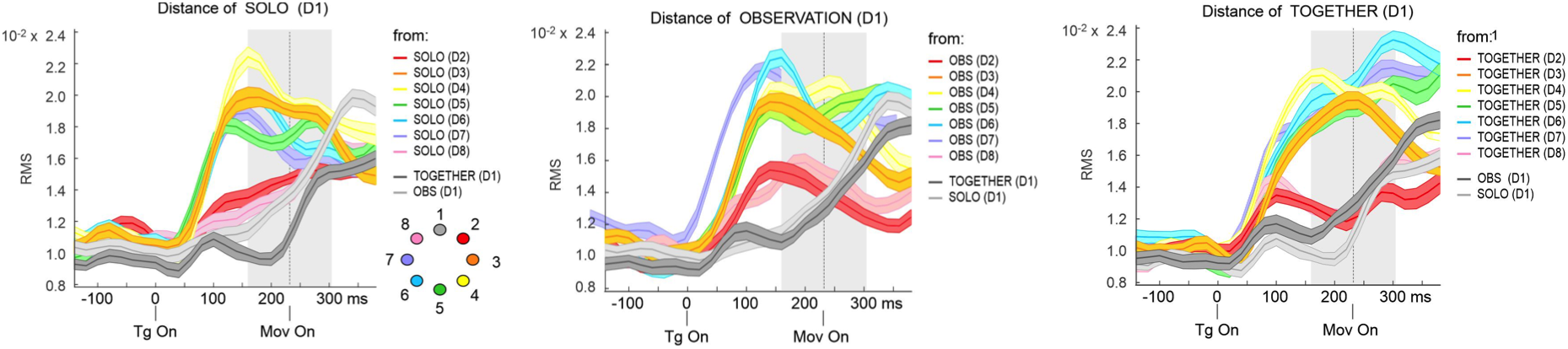
Example of comparisons of neural dynamics across tasks and directions. The comparisons are performed by computing the distance (RMS) in the 12D neural space, between pairs of neural trajetories which differ in the task (gray curves), or in the directions (colors). (***A***) RMS distances between the neural trajectory associated to SOLO condition in direction D1 vs. those differing for task but not direction (OBS-OTHER in D1, light grey, and TOGETHER in D1, dark grey) and for direction but not task (SOLO in D2, SOLO in D3, etc.). (***B-C***) The same as in A, but showing the RMD distances from TOGETHER in D1 (***B***) and OBS-OTHER in D1 (***C***). In all instances, the alignment (0 ms) of activity is to target onset (TgOn). The dotted vertical line and grey shaded area indicate the mean and variance (+/-SD) values of cursor’s motion onset, respectively.

**Figure S8.**
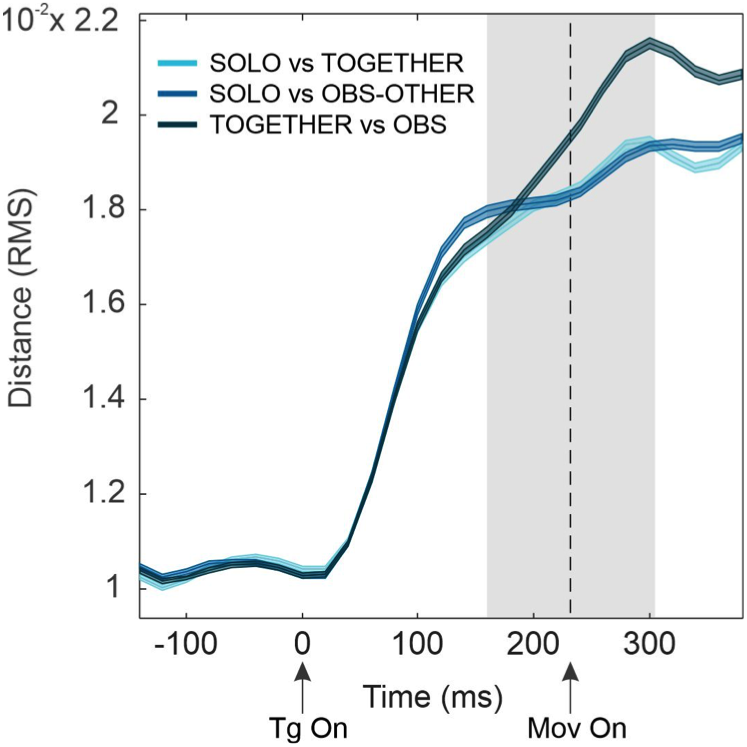
Distances (RMS) in 12D neural space between tasks. The three comparisons (SOLO vs. TOGETHER, SOLO vs. OBS-OTHER and TOGETHER vs. OBS-OTHER) are performed by firstly averaging all the neural trajectories belonging to the same task and then computing their distances (RMS) in the 12D neural space. Alignment (0 ms) is to target onset (TgOn). The dotted vertical line and grey zone indicate the mean and variance (+/-SD) values of cursor’s motion onset, respectively.

